# Prior exposure to malaria decreases SARS-CoV-2 mediated mortality in K18-hACE2 mice without influencing viral load in lungs

**DOI:** 10.1101/2024.02.19.579434

**Authors:** Shradha Mawatwal, Amruta Mohapatra, Sayani Das, Aisurya Ray, Ratnadeep Mukherjee, Gulam Syed, Shantibushan Senapati, Balachandran Ravindran

## Abstract

**Background:** Epidemiological evidence for decreased prevalence and/or mortality due to SARS-CoV-2 infections in countries endemic for malaria have been reported. However, such associational studies in human population are limited by known and several unknown confounding factors. The current study, the first of its kind, was designed to seek experimental evidence to test the hypothesis if prior exposure to Plasmodial infections cross-protect against SARS-CoV-2 challenge infection in a murine model, K-18 human ACE2 transgenic mice.

**Methods:** Mice that had recovered from *<u>Plasmodium chabaudi</u>* infection 40 days earlier were challenged with a virulent strain of SARS-CoV-2 and viral load in lungs as well as mortality were scored and compared with <u>K18 hACE2</u> mice that had not experienced prior malaria.

**Results:** The viral load in lungs 6 days post challenge were comparable in malaria recovered mice and controls suggesting no significant generation of anti-viral immunity. However, mice with prior malaria exposure were significantly protected against SARS-CoV-2 induced mortality. Significant differences were observed in several host immune responses between the two groups when cytokines, chemokines and transcription factors were quantified in lungs. The plasma levels of several cytokines and chemokines were also significantly different between the two groups.

**Conclusion:** The results of the study suggest that prior exposure to malaria protects mice against viral induced mortality in K18 hACE2 transgenic mice challenged with a virulent isolate of SARS-CoV-2 in the absence of demonstrable host immunity inhibiting viral growth in lungs.

## Introduction

The geographical distribution of the pandemic due to COVID-19 has been variable. Infection rate as well as severity and mortality have been found to be generally less in developing economies in comparison to developed countries even after accounting for decreased level of testing and mortality reporting (1). Significant inverse relationship between prevalence of malaria and COVID-19 infection as well as severity and mortality have been reported suggesting possible cross-protection by malaria against the viral disease [2, 3]. While some of these reports are speculative [3–5], other potential reasons for the observed association have also been proposed. Cross reactivity of malarial antibodies with SARS-CoV-2 viral components [2, 6] and differential host immune responses that are functions of plasmablasts and atypical memory B cells generated by malaria have been proposed as possible mechanism for the observed \inverse association [7] including cross reactivity of malarial antibodies to SARS-CoV-2 epitopes [8, 9]. Although these reports are suggestive of possible cross-protection rendered by past exposure to malaria to COVID-19, unequivocal direct evidence has not been reported in literature so far to support the hypothesis.

We proposed that testing in a suitable animal model which has susceptibility to both malaria and SARS-CoV-2 infections could offer direct and more definitive empirical evidence. Two models of SARS infections have been widely used by investigators although neither of them truly represent SARS-CoV-2 infections observed in infected humans - a hamster model in which intranasal exposure of animals to SARS-CoV-2 viral challenge results in demonstrable pathology and viral load in lungs [10] and a mouse model transgenic for human ACE-2 receptor on intranasal exposure to virulent isolates of SARS-CoV-2 displaying very high viral load in lungs leading to mortality of about 85-90% of infected animals has been used extensively for testing drugs and vaccines [11, 12]. While hamsters are not suited to study experimental malarial infections, a model of murine malaria that experiences an acute self-limiting parasitaemia has been used in the current study (13). In this model mice sustain microscopically undetectable low grade parasitaemia for several months after experiencing a course of acute infection followed by microscopically undetectable parasitemia for several months. This model also mimics the asymptomatic malarial infections observed in many subjects living in disease endemic countries. Use of such a recovering murine model of malaria allowed us to challenge them with a virulent SARS-CoV-2 isolate in K-18 h ACE2 transgenic mice to address the issue of malaria offering cross protection to SARS CoV-2 infections.

We challenged K18 hACE2 transgenic mice with a virulent strain of SARS-CoV-2 viral isolate 4-5 weeks post recovery from acute *<u>Plasmodium chabaudi</u>* infection and scored for viral load and activation of several host genes in lungs 6 days post infection. Another cohort of mice were challenged and followed over a period of 35 days to score mortality and hypothermia. The investigations provided experimental evidence to suggesting absence of cross protecting anti-viral immunity to SARS CoV-2 infection by malaria, although prior exposure to malaria altered host immune responses and significantly protected mice from SARS-CoV-2 induced mortality.

## Methods

### Animals

B6.Cg-Tg(K18-ACE2)2Prlmn/J Strain #:034860 (common name: K18-hACE2) mice imported from Jackson laboratory, USA were used in in-house ABSL (animal bio-safety level 3 laboratory) facility – details of mice strain is available at https://www.jax.org/strain/034860. The animals were bred in the ILS central animal facility and were genotyped for human ACE2 transgene and hemizygotes were used for all infections. The K18-Human ACE2 transgenic mouse model recapitulates non-severe and severe COVID-19 in response to an infectious dose of the SARS-CoV-2 virus

### Ethics and bio-safety approval

The project “Immunomodulation in chronic murine malaria” was approved by Institutional Animal Ethics Committee and was conducted as per approved guidelines. Institutional bio- safety committee approved all experimental protocol for viral infection and were performed in biosafety level 3 laboratory as described below.

### SARS CoV-2 strain and infection

A SARS-CoV-2 strain of clade 19A, IND- ILS 01/2020 (Genbank accession ID- MW559533.2) isolated from a clinical sample in Odisha, India during the 2019 pandemic was used in the study [10, 14]. The virus was propagated in Vero-E6 cells in DMEM containing 2% FBS supplemented with antibiotics and the end-point infectivity titre (plaque-forming unit) was determined as described [15]. All experiments with the SARS- CoV-2 were performed in the level 3 containment facility (BSL-3) of the Institute of Life Science (ILS), Bhubaneswar, India as per protocol approved by the Institutional Biosafety Committee (IBSC) guidelines.

### *Plasmodium chabaudi* adami strain and infection

The strain of *Plasmodium chabaudi adami* was received from Prof. Shobona Sharma, Tata Institute of Fundamental Research, Mumbai [13] and blood collected from infected C57 Black6 mice were stored in 70% glycerol. The parasites were revived by injecting in normal mice which were then used as donors to infect <u>K18 hACE2</u> and wild type C57BL/6 mice by intraperitoneal route with a dose of 5x 10^5^ parasitized erythrocytes. The parasitemia were scored by microscopic examination of Giemsa-stained thin blood smears. A characteristic self-limiting infection was observed in both the strains of mice with parasitemia reaching about 20- 35%. The <u>K18 hACE2</u> mice suffered significantly higher parasitemia in comparison to B-6 mice (Fig S4a) The mice were free of microscopically detectable parasite in thin blood smears after 30 days although presence of parasites could be detected in peripheral blood by PCR for several months post recovery (Fig 3d)

### Experimental design

The study was conducted by following two protocols of SARS-CoV-2 infection

### Acute SARS-CoV-2 infection model

Two groups of K18 hACE2 mice were infected with SARS-CoV-2 by intranasal route – one that had previously recovered from *Plasmodium chabaudi* infection and the other control groups of mice. The animals were euthanized on day 6 post challenge and lungs were collected for analysis. Blood collected in EDTA were used for separating plasma samples. <u>The schematic</u> <u>diagram in</u> Fig 1a shows the broad design of the experiment and the assays performed. Details of each of the assays is described below.

**Fig 1:**
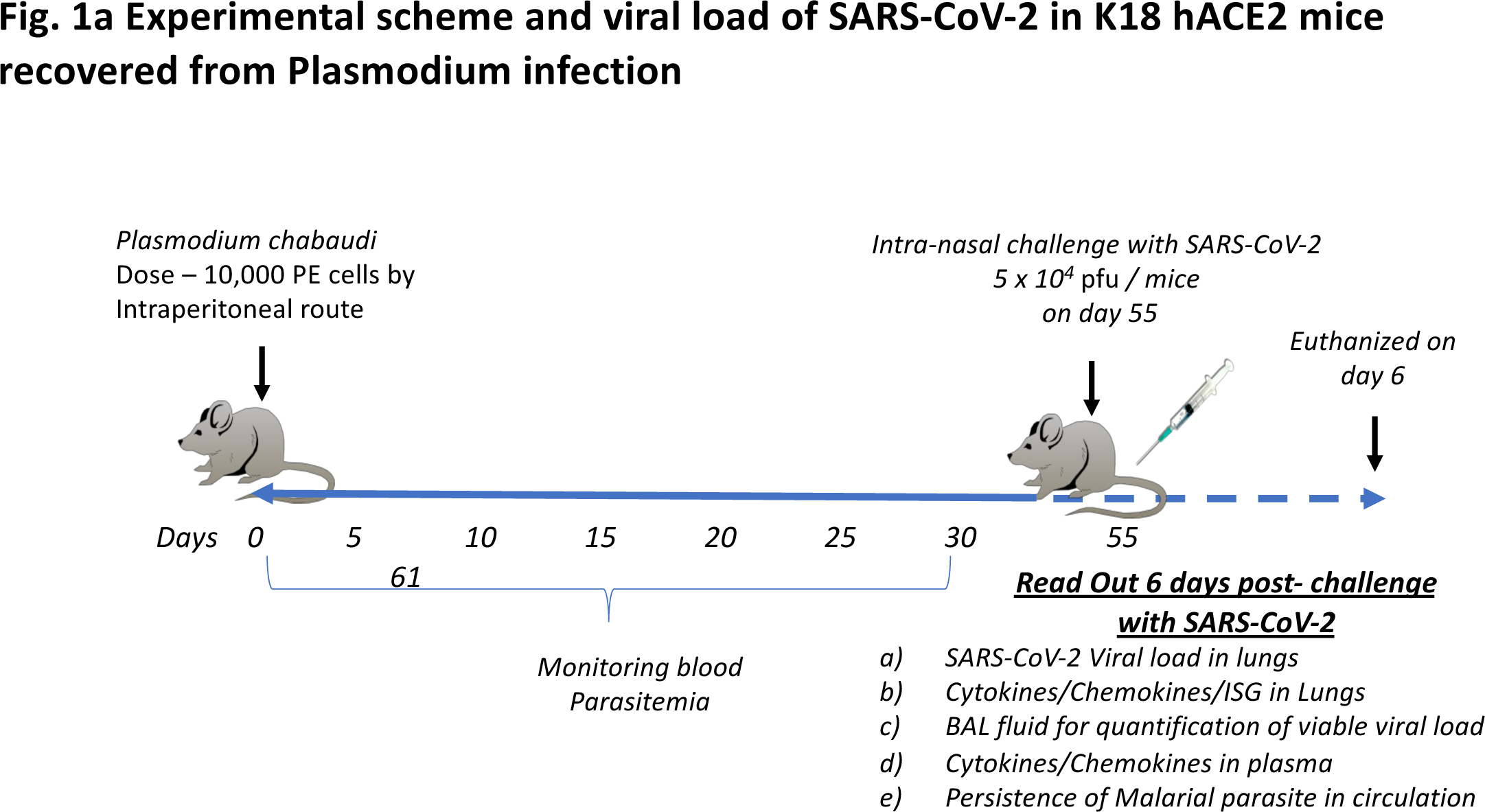
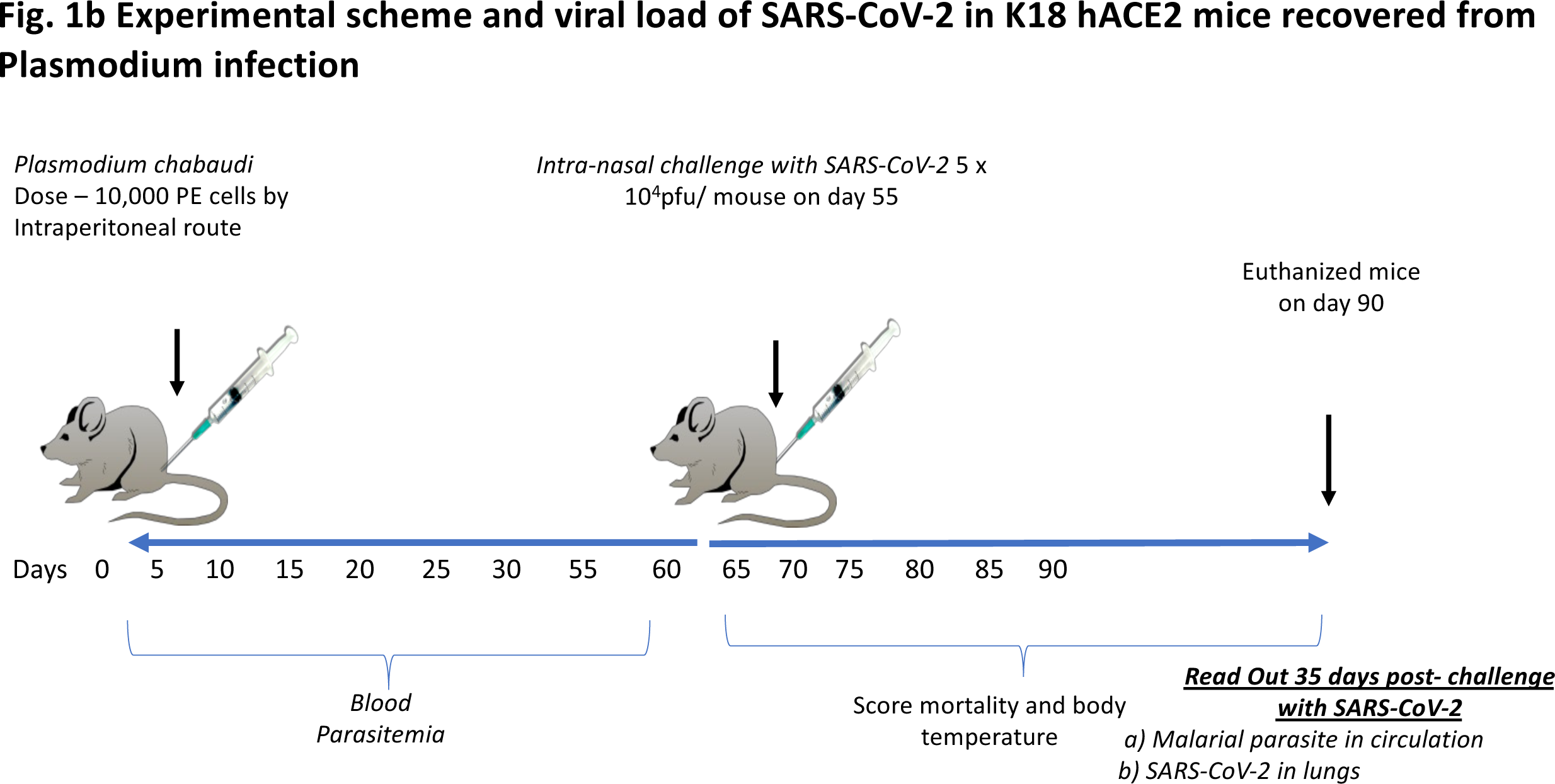
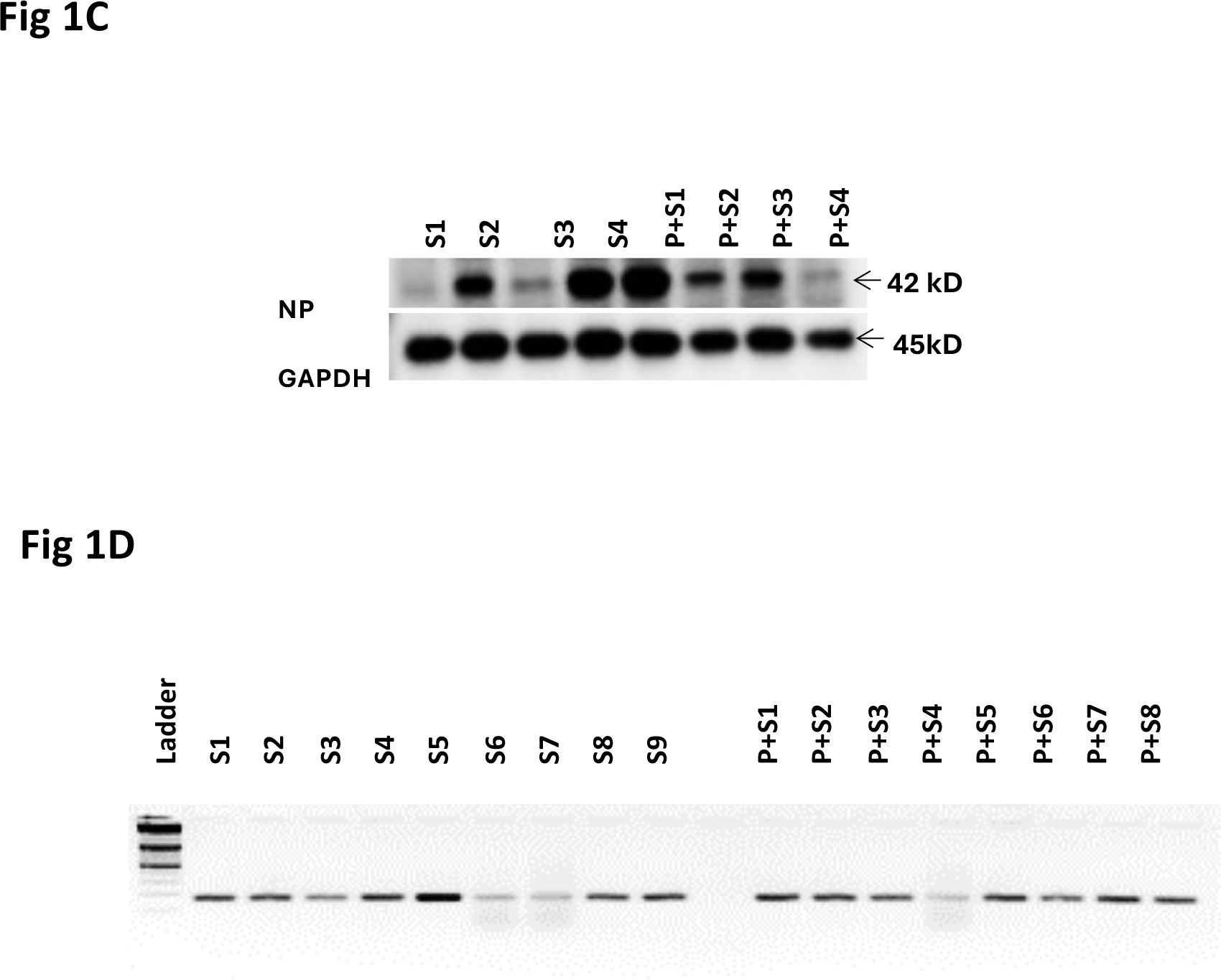
Experimental scheme and viral load of SARS-CoV-2 in control and Plasmodium chaubudi recovered K18-hACE2 mice. **Fig 1a) Experimental scheme of K18-hACE2 acute mouse model of SARS-CoV-2 :** K18- hACE2 mouse infection model. Mice (N=10) were intraperitonially infected with *Plasmodium chabaudi* at a dose of 10^4^ parasitized erythrocytes followed by monitoring parasitaemia in circulation in Giemsa stained thin blood smears. . The animals recovered from microscopically demonstrable parasitaemia were rested. On 55^th^ day, mice were intra-nasally challenged with SARS- CoV-2 at 5×10^4^ PFU/mice along with a control group of K18-hACE2 mice (N=10). Blood, bronchial lavage (BAL) and lungs were collected at 6 days post infection for analysis as shown in the cartoon. **Fig 1b) Experimental scheme of K18-hACE2 chronic mouse model of SARS-CoV-2 :** K18- hACE2 mice (N= 9) were intraperitonially infected with *Plasmodium chabaudi* at dose of10^4^ parasitized erythrocytes followed by monitoring parasitaemia in circulation. The animals recovered from microscopically demonstrable parasitaemia and were rested. On 55^th^ day, the mice were intra-nasally infected with SARS- CoV-2 at 5×10^4^ PFU/mice along with a control group of K18-hACE2 mice (N=7). The experimental and control groups were monitored for mortality and body temperature. The animals were euthanized 35 days post challenge with SARS-CoV-2 and blood and lungs were collected for scoring persistence of malarial parasite and SARS-CoV-2 virus. Fig 1c) Western blot analysis for nucleocapsid protein expression in representative lung lysates (6 days post challenge) of K18 hACE2 mice infected only with SARS- CoV-2 (S1 to S4) and 4 mice infected prior with Plasmodium (P+S1 to P+S4) followed by challenge with SARS- CoV-2. Fig 1d) PCR for SARS-CoV-2 nucleocapsid gene expression in c-DNA of lung tissues (6 days post challenge) of K18 hACE2 mice infected only SARS-CoV-2 (S1 to S9) and mice infected prior with Plasmodium (P+S1 to P+S8) followed by challenge with SARS-CoV-2.

### Chronic SARS-CoV2 infection model

Two groups of <u>K18 hACE2</u> mice were infected with SARS-CoV-2 by intranasal route – one that had previously recovered from *<u>Plasmodium chabaudi</u>* infection and the other control groups of mice. The animals were followed up for a period of 35 days and mortality and body temperature were recorded analysed as described in Fig 1b. On day 35 post challenge the animals were euthanized and lungs were collected for analysis. Blood collected in EDTA were used for separating plasma samples. The <u>schematic diagram in</u> Fig 1b shows the broad design of the experiment and the assays performed. Details of each of the assays is shown described below.

### PCR and qRT-PCR for SARS CoV-2 viral load in lungs

Mouse whole lung tissues were weighed and homogenized. Total RNA was extracted from mouse tissue samples using Trizol LS reagent (Thermo Fisher Scientific, IL, USA). First-strand cDNA synthesis of Isolated RNA was carried out using random primers and high capacity cDNA reverse transcription kit (Thermo Fisher Scientific, IL, USA). The forward and reverse primers used for SARS-CoV-2 nucleocapsid were 5’-GTAACACAAGCTTTCGGCAG-3’ and 5’-GTGTGACTTCCATGCCAATG-3’ respectively. Real-time quantitative PCR was performed using SYBR select master mix (Thermo Fisher Scientific, IL, USA). The reaction conditions were as follows: 95 °C for 30 sec, followed by 40 cycles of denaturation at 95 °C for 5 sec and extension at 58 °C for 34 sec in the QuantStudio 6 system (Applied Biosystems Co., USA). At the end of PCR, dissociation curve analysis was performed to verify the amplification of a single product. The standard curve for absolute quantification of viral genome copies was generated using log-fold dilutions of plasmid harbouring the SARS-CoV- 2 nucleocapsid gene. PCR was performed in 20 μl of PCR solution containing 50 mM KCl, 10 mM Tris HCl (pH 8.3), 2.5 mM MgCl2, 200mM (each) dNTP, 50 pmol of each primer, 0.1 mg of gelatin per ml, 1U of Taq DNA polymerase (Promega, Leiden, The Netherlands). Amplification was accomplished over 30 cycles as follows: Reaction mixtures were pre-incubated for 5 min at 95°C for DNA denaturation followed by heating to 95°C for 30 sec prior to cycling at 60°C for 30 sec of annealing, 72°C for 1 min of extension, and 72°C for 10 min of final extension. The PCR products were analysed by 2% agarose electrophoresis.

### PCR for *Plasmodium chabaudi* in mice blood

At 35 days after SARS infection blood was collected in EDTA-coated tubes by cardiac puncture. DNA from the blood of *Plasmodium chabaudi* infected mice was isolated from QIAmp DNA blood kit (QIAGEN, Valencia, CA) as per the manufacturer instruction. Briefly, protease k was added to blood followed by the lysis buffer. The blood was incubated for 10 min at 56°C followed by addition of absolute ethanol. After washing the DNA was eluted in elution buffer. The samples were screened for Plasmodium using a two-step PCR procedure developed by Perkin et al. 1998 [16]. In the first reaction, genus-specific oligonucleotide primers from Li et al. 1995 (5’-CGACTTCTCCTTCCTTTAAAAGATAGG-3’ and 5’- GGATAACTACGGAAAAGCTGTAGC-3’) were used to amplify an approximately 1200-bp region of the 18S Plasmodium ribosomal sub-unit gene. PCR was performed in 20 μl of PCR solution containing 50 mM KCl, 10 mM Tris HCl (pH 8.3), 2.5 mM MgCl2, 200mM (each) dNTP, 50 pmol of each primer, 0.1 mg of gelatin per ml, 1U of Taq DNA polymerase (Promega, Leiden, The Netherlands). Amplification was accomplished over 35 cycles as follows: Reaction mixtures were pre-incubated for 5 min at 95°C for DNA denaturation followed by heating to 95°C for 60 sec prior to cycling at 48°C for 60 sec of annealing, 72°C for 2 min of extension. The second reaction utilized the nested primers 5’- TAACACAAGGAAGTTTAAGGC-3’ and 5’- TATTGATAAAGATTACCTA-3’ (Li et al. 1995). Each 20-µl reaction again received 1µl of each primer and PCR solution as well as 1µl of product from the first reaction, serving as target DNA. Thermocycling conditions were identical to those in the first reaction, except that the annealing temperature was raised to 50°C. The products were run out on an acrylamide gel, stained with ethidium bromide, and were scored positive if a band of approximately 420-bp, indicative of the 18S Plasmodium ribosomal sub-unit gene, was apparent.

### qRT-PCR for cytokines and IFN-β

Total RNA was extracted and cDNA was prepared as described above. qRT-PCR was performed to quantify the following cytokine genes and IFN-β in lung tissue. Real-time quantitative PCR was performed using SYBR select master mix (Thermo Fisher Scientific, IL, USA). The reaction conditions were as follows: 95 °C for 5 min, followed by 40 cycles of denaturation at 95 °C for 30 sec and extension at 60 °C for 30 sec in the QuantStudio 6 system (Applied Biosystems Co., USA). At the end of PCR, dissociation curve analysis was performed to verify the amplification of a single product. For assay normalization, the housekeeping gene β-actin was used. The fold-change in expression was calculated by the 2^−ΔΔCt^ method. The following gene-specific primer sequences for cytokines, and IFN-β was used for qRT PCR:

**Table 1:**
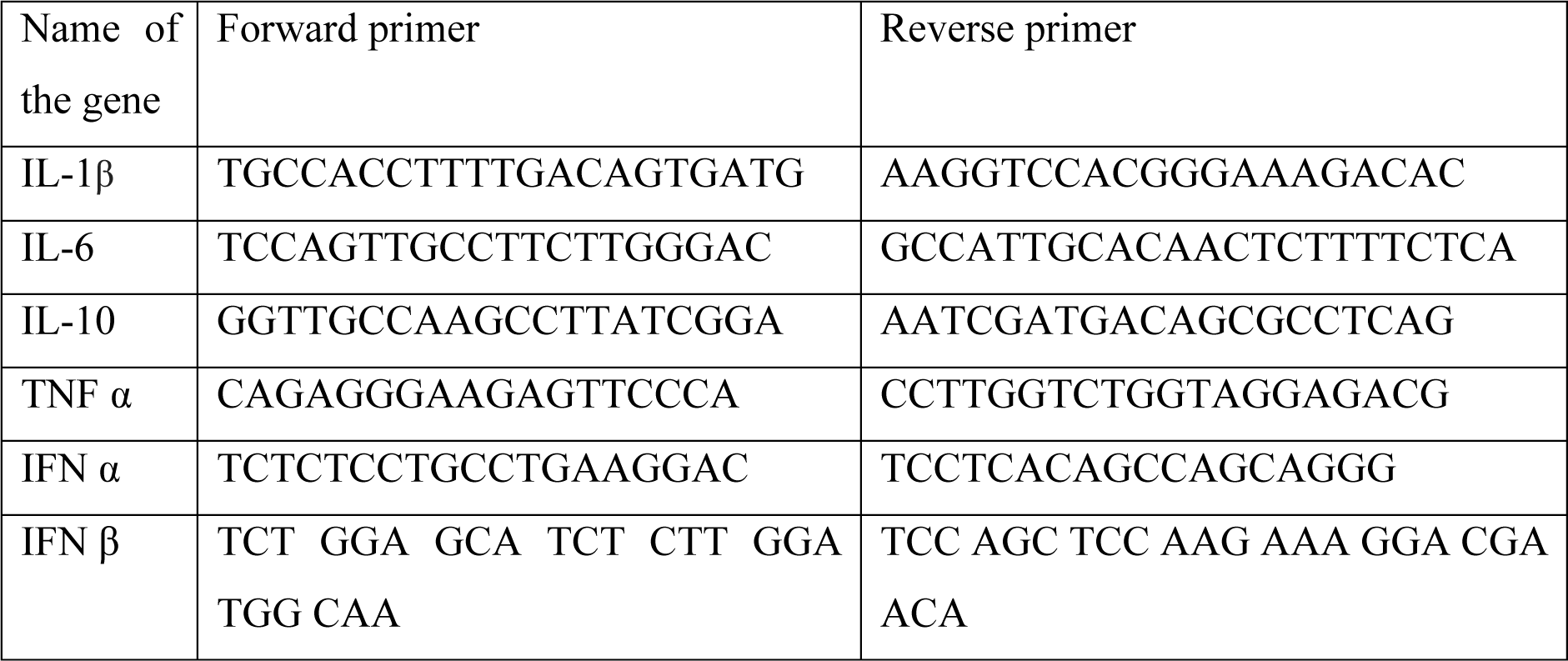
qRT-PCR primer list for cytokines and IFN subtypes.

### ELISA for type 1 Interferon

The level of IFN- β was determined in mice plasma by solid phase sandwich ELISA using Abcam IFN- β ELISA kit (Burlingame, CA, USA) in accordance with manufacturer’s instructions. Concentration of the cytokine was estimated by testing standards and plasma followed by addition of specific capture antibody- enzyme conjugate to the wells. After 1 hr incubation at room temperature, the plates were washed followed by addition of TMB development solution and read within 20 min of stopping reaction at 450 nm in a plate reader. Concentration of the cytokine in culture supernatant was measured through linear regression analysis using standard curve as per manufacturer’s instructions and expressed as pg/ml.

### qRT-PCR for ISGs and Chemokines in lungs

RNA was isolated from lung homogenates and c-DNA were prepared as described above. Real-time quantitative PCR was performed using SYBR select master mix (Thermo Fisher Scientific, IL, USA). Amplification was accomplished over 30 cycles as follows: 95 °C for 15 sec and 60 °C for 1 min. At the end of PCR, dissociation curve analysis was performed to verify the amplification of a single product. For assay normalization, the housekeeping gene β-actin was used. The fold-change in expression was calculated by the 2 ^−ΔΔCt^ method. The following gene-specific primers were used:

**Table 2:**
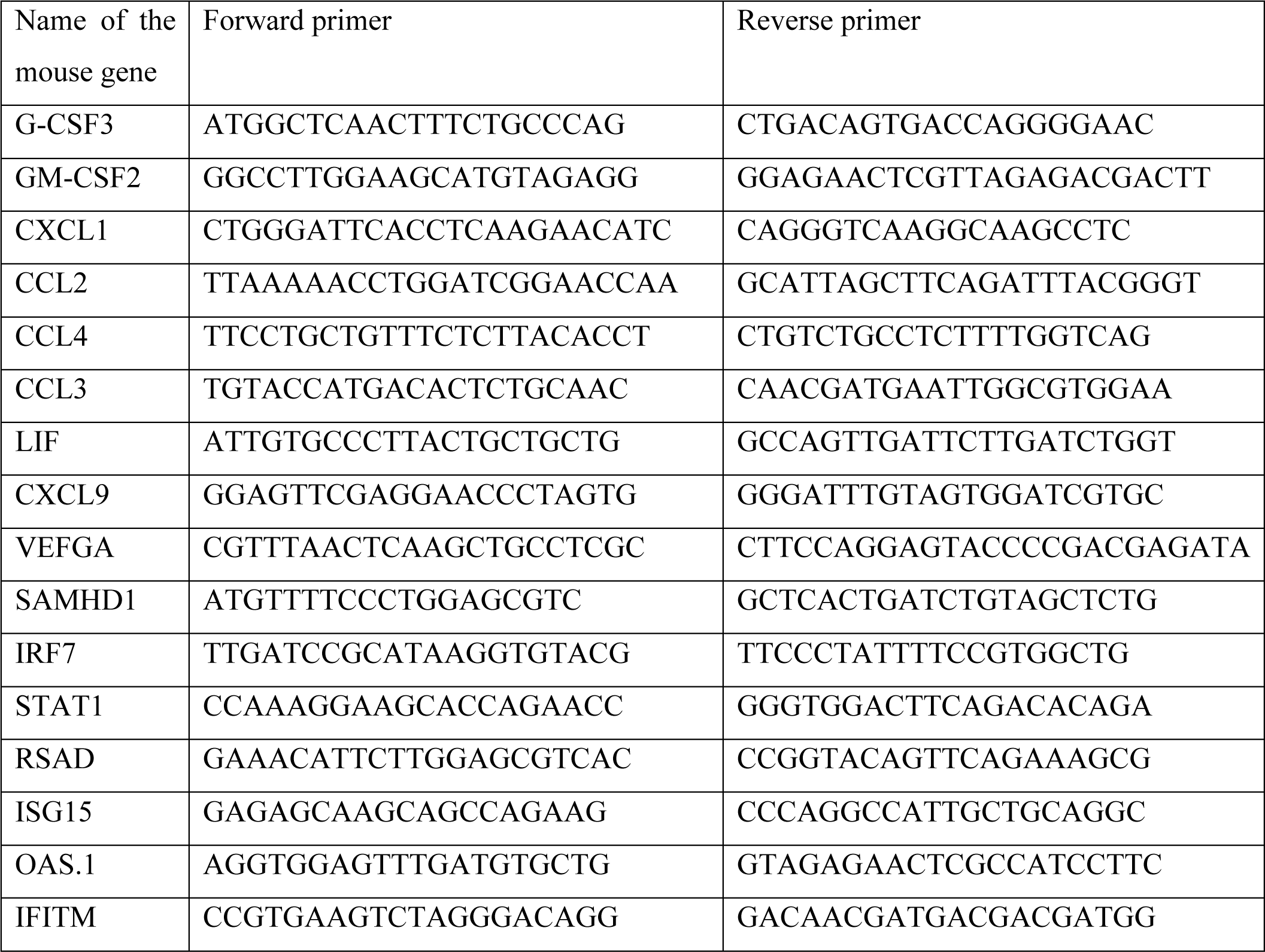
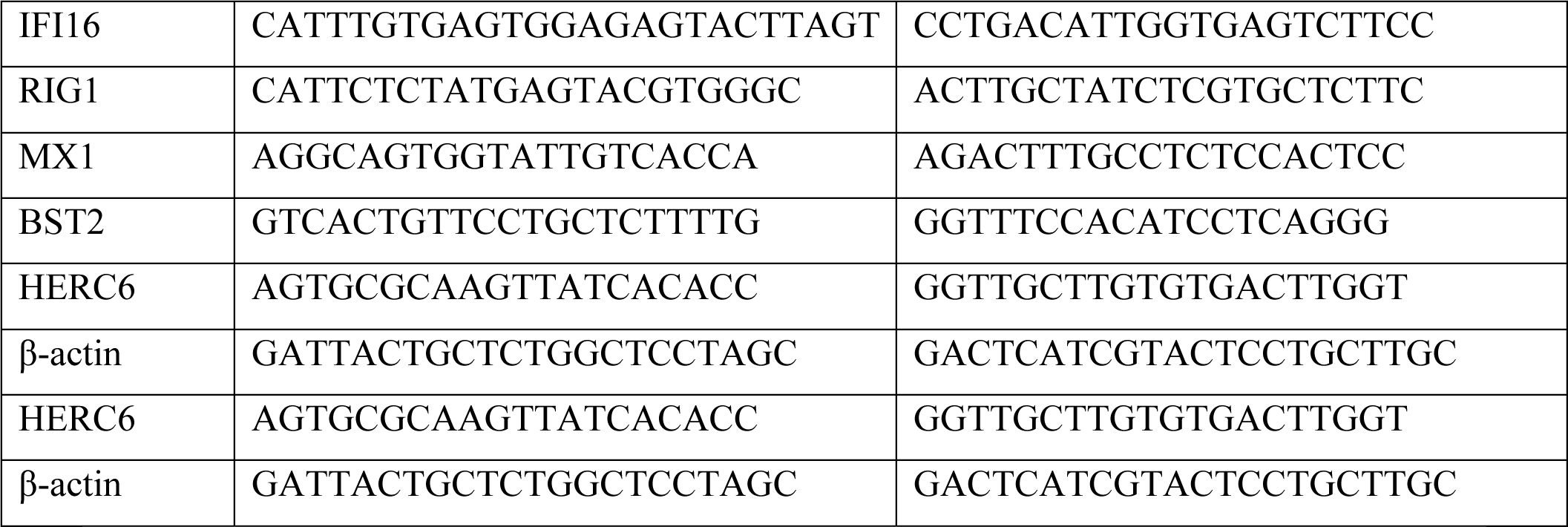
qRT-PCR primer list for ISG and chemokines.

### Western blotting

Lung tissue samples were homogenized in the radio-immunoprecipitation assay (RIPA) buffer (20 mM Tris, pH 8.0; 0.1% Sodium deoxycholate; 1% Triton X 100; 0.1% Sodium Dodecyl Sulphate; 150 mM NaCl) containing protease inhibitor cocktail using an electric homogenizer. Tissue lysates were separated by centrifugation and separated by SDS-PAGE electrophoresis and transferred onto a nitrocellulose membrane (Bio-Rad). The membrane was blocked for 1 hr in 5% skimmed milk followed by incubation with primary antibody specific for SARS-CoV- 2 nucleocapsid protein (1:300; Abgenex #11-2003) overnight at 4°C. The membrane was then washed thrice with 1X PBST and incubated for 1 hr with HRP conjugated secondary antibody (1:5000; Promega #w4018). After washing with PBST the membrane was developed using the enhanced chemiluminescence reagent (Bio-Rad Laboratories, Hercules, CA).

### Bead assay for cytokines and chemokines

Bioplex for cytokines/chemokines <u>(23-plex)</u> for quantitative estimation of the following molecules: IL-1α, IL-1β, IL-2 (36), IL-3 (18), IL-4 (39), IL-5 (52), IL-6 (38), IL-9 (33), IL-10 (56), IL-12p40 (76), IL-12p70 (78), IL-13 (37), IL-17A (72), Eotaxin (74), G-CSF (54), GM-CSF (73), IFN-γ (34), KC (57), MCP-1 (51), MIP-1a (77), MIP-1b (75), RANTES (55), TNF-a (21). Univariate analysis was performed for comparing each of the molecules between the two groups. Principal components analysis (PCA) was used for the dimensionality reduction of multivariate cytokine responses. log(n+1) values of cytokine concentrations were used for PCA and the samples were visualized on a two-dimensional plot of the first two principal components. A factor loading plot to visualize the contributions of individual cytokines toward the principal components was also undertaken. PCA was performed with the scikit-learn package in Python (v 3.9.13)

### Immunofluorescence assay for quantifying antibodies to SARS-CoV-2

Vero E6 cells were seeded at 70% confluency in 96-well black optical bottom plates. After 12 hr, the cells were infected with 1 MOI of SARS-CoV2 virus diluted in 2% FBS-containing culture medium for 2 hr at 37°C. The virus inoculum was then replaced with 10% FBS-containing culture media. After 36 hr post-infection, the cells were washed with PBS and fixed in 4% paraformaldehyde. Subsequently, the cells were permeabilized and blocked for 1 hr with PBS containing 5% BSA and 0.1% TritonX-100. Later, the cells were incubated overnight at 4°C with pre-clarified plasma obtained from mice challenged with respective pathogens at 1:100 dilution in antibody binding buffer. A commercial immunofluorescence grade primary antibody specific for SARS-CoV2 Spike RBD domain (Abgenex, Cat. No. 10-10036) was used as a positive control for infection. After primary sera incubation, the cells were washed 3x with PBS for 5 mins each, followed by incubation with Alexa Fluor 568 conjugated donkey anti-mouse secondary antibody (Invitrogen) for 1 hr at room temperature in the dark. After 3x washes with PBS for 5 mins each, fluorescence reading was taken in Varioskan Flash, Multimode reader (Thermo Scientific) with 578 nm and 603 nm excitation and emission setting. The relative fluorescence intensity observed in the corresponding mock- and SARS- CoV2-infected Vero E6 cells incubated with respective plasma was plotted by subtracting the only secondary antibody control values from the observed fluorescence intensity.

### Foci Forming Unit Assay (FFA) in bronchial fluid

Vero cells were seeded in 96 well black optical bottom plates at 70% cell density (Nunc- immunoTM). After 12h, the cells were infected with 10 fold serial dilutions of Broncho- alveolar lavage (BAL) fluid for 2 h at 37°C in a humidified 5% CO2 incubator. After 36 h incubation, cells were washed with 1X PBS and fixed in 4% paraformaldehyde. Subsequently the cells were permeabilized and blocked for 1h with PBS containing 3% BSA and 0.1% TritonX-100, followed by incubation with primary antibody specific for SARS-CoV2 Spike RBD domain (Abgenex, Cat. No. 10-10036) overnight at 4°C. After 3 washes with 1X PBS, the cells were probed with donkey anti-mouse secondary antibody tagged with Alexa Fluor 568 (Invitrogen), for 1 h in dark, at room temperature. After 4 washes with 1x PBS, the cells were counter stained with DAPI for 10 min. Images were captured under a 10x objective using an Olympus DX58 inverted fluorescence microscope. The infection foci were counted and virus titres calculated as FFU/mL culture supernatant (Singh, Avula et al. 2022).

### Statistics

Statistical tests were performed with GraphPad Prism, 5.0 version (GraphPad Software, San Diego, CA). Data are presented as mean ± SEM. Comparison of 2 groups was performed using Student’s two-tailed t test (for unpaired samples) or a paired t test (for paired samples). The *p* values were assessed using two-tailed unpaired or paired Student’s t test with *p* values considered significant as follows: \**P* < 0.05, \*\**P* < 0.01, \*\*\**P* < 0.001.

## Results and discussion

### Effect of prior malaria exposure in acute model of SARS-CoV-2 challenge infection in K18 hACE2 mice

Mice infected with *<u>Plasmodium chabaudi</u>* after a phase of self-limiting parasitaemia recover and the parasites persist at submicroscopic levels. Significant hematological changes such as enhanced granulocytes and mononuclear cells and higher anemia were observed in K18 mice in comparison to wild type C57BL/6 mice (Fig S4 a-e). All the hematological changes returned to normal levels around 30 days post infection. K18 mice were significantly more susceptible to malarial parasitemia and were comp[arable to wild type mice in terms of recovery from malaria. About 40 days post recovery from malaria the animals were infected by intranasal exposure to SARS-CoV-2 and 6 days post infection, the viral load in lungs were analysed by PCR <u>and qRT-PCR</u> (Fig 1c, 2a respectively) and by western blot (Fig 1d). There were no significant differences between the two groups. Malaria pre-infected and challenged with SARS CoV-2 and the control group infected only with SARS-CoV-2 were comparable suggesting absence of direct anti-viral cross protection by malaria towards SARS-CoV-2 at the time point of 6 days post challenge (Fig 1c, d, Fig 2a). These observations in experimental and control mice were further confirmed in a Foci forming unit assay using bronchial fluid collected 6 days after challenge with SARS-CoV-2. (Fig 2b). This assay allows for measurement of viable virus titer in biological samples [15, 17 and 19]. However, profiling cytokine in the plasma (collected on day 6) of mice infected either with SARS-CoV-2 or pre-infected with malaria followed by SARS CoV-2 challenge revealed significant differences in host response. Univariate statistical analysis revealed plasma levels of IL-9, IL-10, IL-12p40, Eotaxin, GM-CSF, KC, and MCP-1 to be significantly low in animals with prior exposure to malaria in comparison to controls (Fig 2c) and no significant differences in all other cytokines/chemokine levels were observed. It is worth noting that the significantly different cytokines have distinct functional profiles: GM-CSF, IL-9, KC (CXCL1), and MCP-1 in homeostasis, neutrophil recruitment, and chemotaxis; IL-10 and Eotaxin during anti-inflammatory response and IL- 12p40 suggestive of Th1 differentiation. Lower levels of all the above cytokines in the test group is suggestive of differences in hematopoiesis and cellular homeostasis and inflammatory responses in mice that had experienced malaria and subsequently infected with SARS-CoV-2. A combined cytokine profile that can discriminate the two treatment groups was performed. Principal component analysis (PCA) shows that systemic cytokine response is significantly different between control K18 mice infected with SARS-CoV-2 and SARS-CoV-2 pre-infected with malaria (Figure 2d). This difference was only on the first principal component, which accounts for most of the variability in the data. Inspection of the contribution of individual cytokines toward the first principal component revealed a prominent dichotomy of positive and negative contributors, with IL-6 having the strongest positive contribution, while Eotaxin being the most negative (Fig S1). This suggests that SARS-CoV-2 mice previously infected with malaria have heightened overall host immune response in comparison to mice infected with only SARS-CoV-2. Similar differences were observed when activation profile of genes in lungs for cytokines/chemokines were quantified. Significant down regulation of the following genes was observed in lungs of mice that had experienced malaria earlier: G-CSF, GM-CSF, VEGFA, LIF (Fig 2f). On the other hand, significantly elevated response of several of the Interferon stimulated genes (ISGs) were observed in mice that had experienced malaria earlier (Fig 2e). No significant differences were observed for many of the other cytokines/chemokine and Interferon genes tested (Sup Figs 2a,b,c and d) on day 6 post challenge in both the groups.

**Fig 2.**
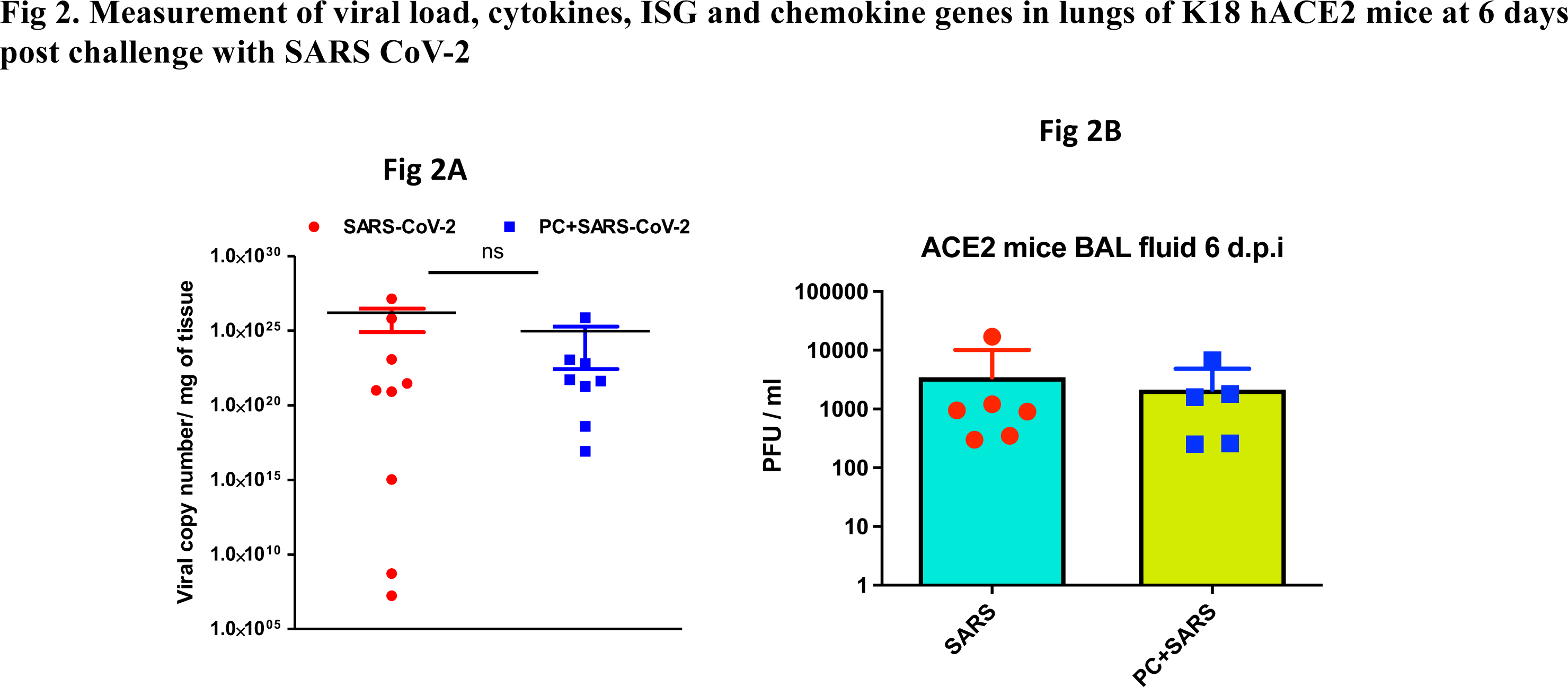
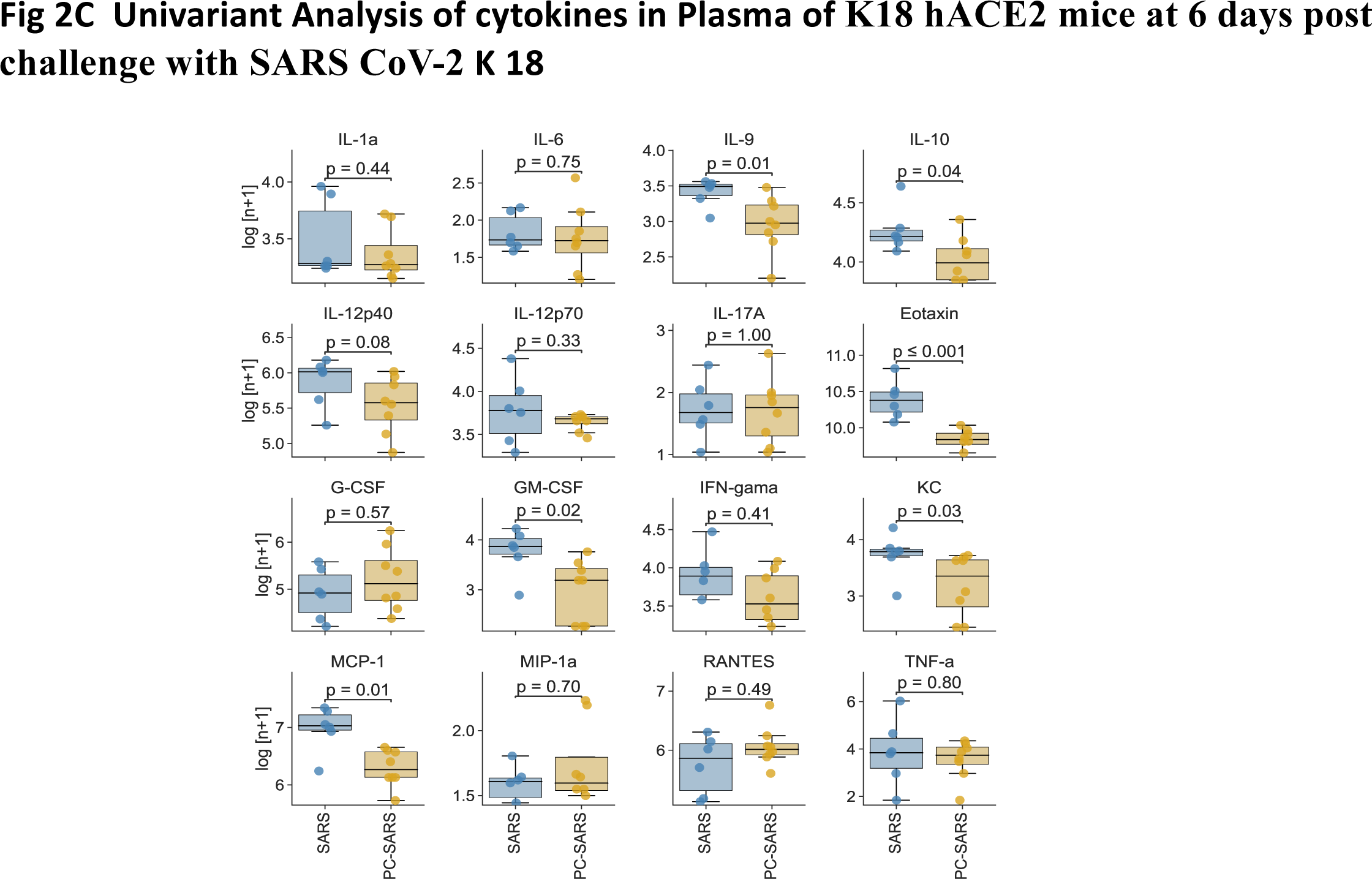
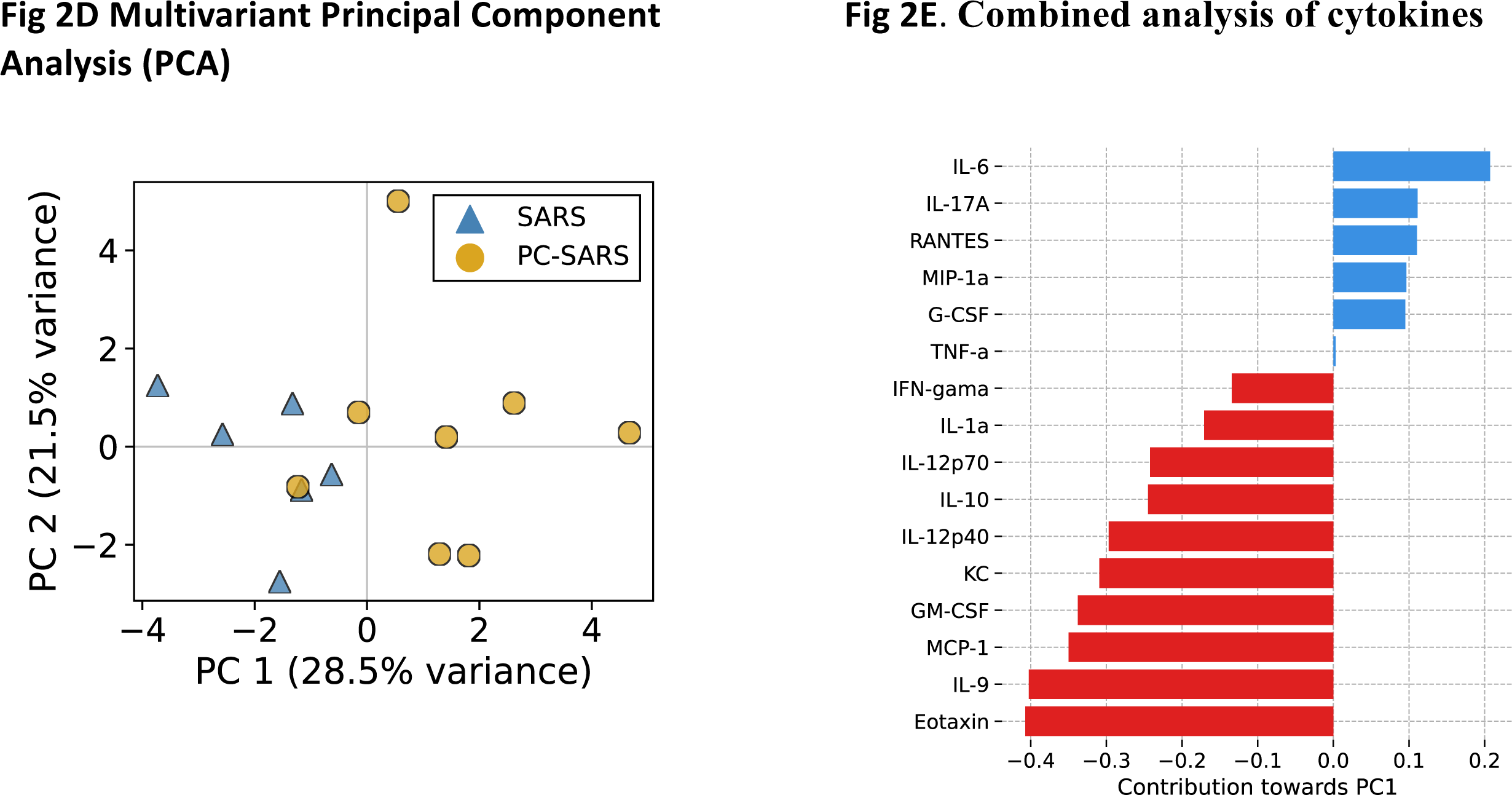
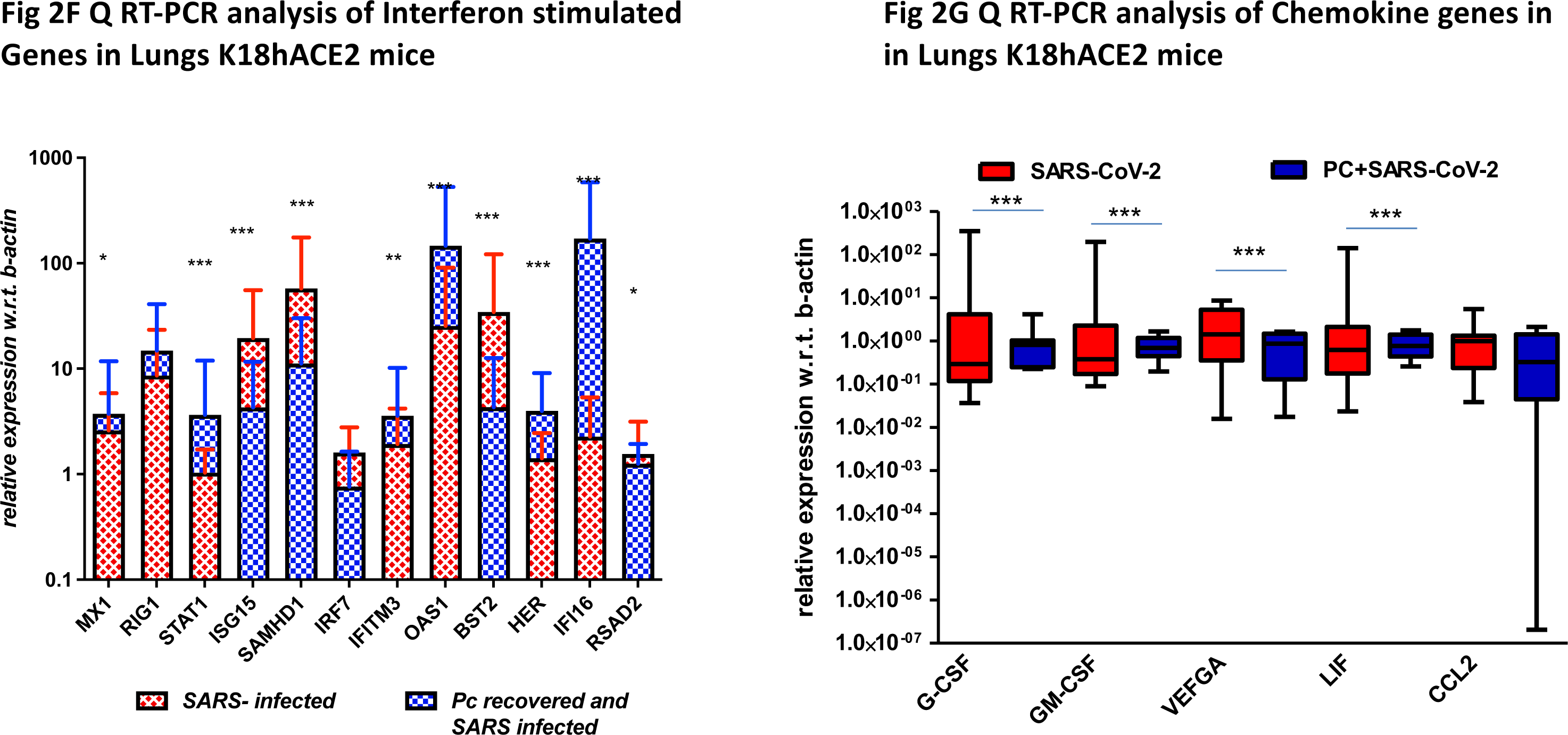
Measurement of viral load, cytokines, ISG and chemokine genes in lungs of K18 hACE2 mice at 6 d.p.i. Fig 2a) RT-PCR for quantification of SARS-CoV-2 viral copy number in lungs of control mice infected with SARS-CoV-2 (N= 9) and in malaria recovered forllwed by challenge with SARS-CoV-2 (N=8). Analysis of lungs 6 d.p.i with the virus by intranasal route is shown. Fig 2b) Foci Forming Unit analysis for quantification of infectious viral titers in lungs of control mice infected with SARS-CoV-2 (n= 6) and mice pre-exposed to malaria and 30 days post-recovery infected with SARS CoV2 (n=5) at 6 d.p.i after SARS-CoV-2 challenge by intranasal route. Data are presented are mean ± SEM of three technical replicates from each individual mouse of the respective groups. Fig 2c) Univariant analysis of 12 of cytokines/chemokines in SARS-CoV-2 infected K18 hACE2 mice (blue) and K18 mice recovered from Malaria followed by challenge with SARS-CoV-2 (deep yellow) is shown as Mean ± SEM for 16 molecules. Plasma samples collected on day 6 post challenge were analysed by bead assay for cytokines/chemokines (Bioplex). Fig 2d) Principal component analysis (PCA): Multivariate analysis of cytokines/chemokines in SARS-CoV-2 infected K18 hACE2 control mice (blue) and K18 hACE2 mice prior exposed to malaria followed by challenge with SARS-CoV-2 (deep yellow) is shown. Plasma samples collected on day 6 post challenge were analysed by bead assay for cytokines/chemokines (Bioplex). Fig 2e. The factor loadings plot for PCA – descriptions same as above shown under Fig2d – Fig 2f) qRT-PCR analysis for indicated ISGs using c-DNA prepared from lungs of control SARS-CoV-2 infected (n = 9) and in mice prior exposed to malaria followed by challenge with SARS-CoV-2 (n = 8) at 6 days post viral challenge. Data are presented as mean ± SEM. Statistical analyses of fold change is shown in asterix performed by Student’s unpaired t-test (*P < 0.05, ***P < 0.0005) between the two groups for each of the ISGs. Fig 2g) qRT-PCR analysis for indicated chemokines using c-DNA from lungs of control SARS-CoV-2 infected (n = 9) and in mice exposed earlier to Malaria followed by challenge with SARS-CoV-2 (n = 8) at 6 days post viral challenge is shown. Data are presented as mean ± SEM of fold changes. Statistical analyses is shown in asterix by Student’s unpaired t-test (*P < 0.05, ***P < 0.0005) between the two groups for each of the ISGs.

**Fig 3:**
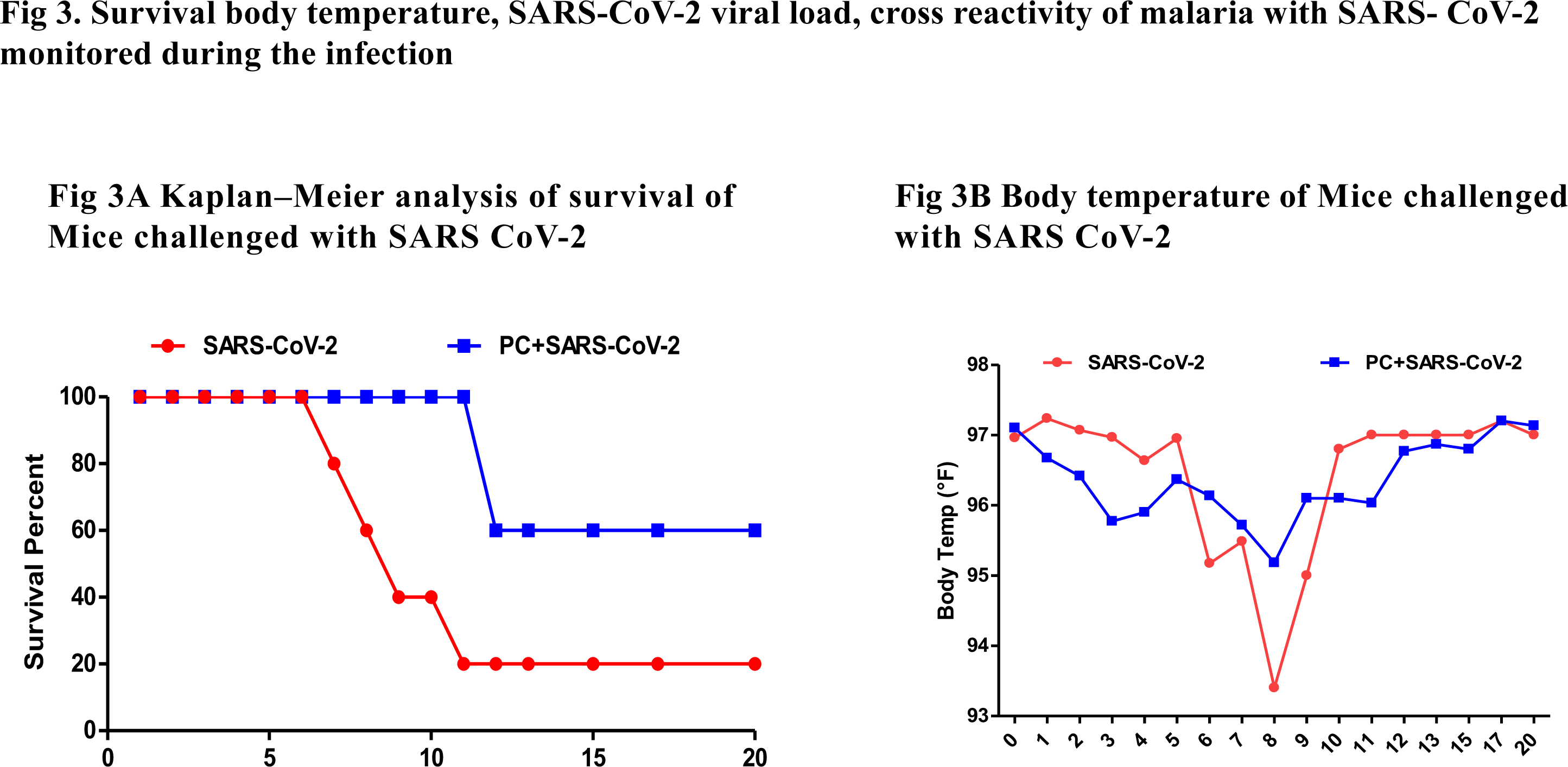
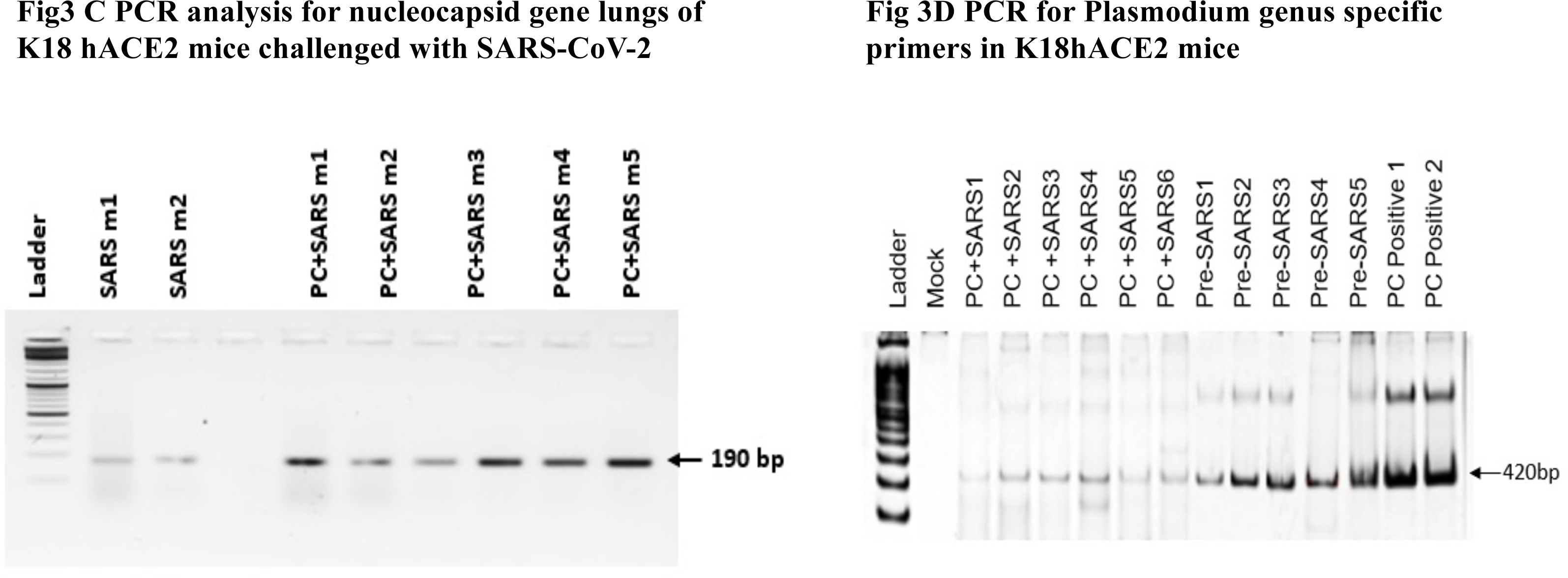
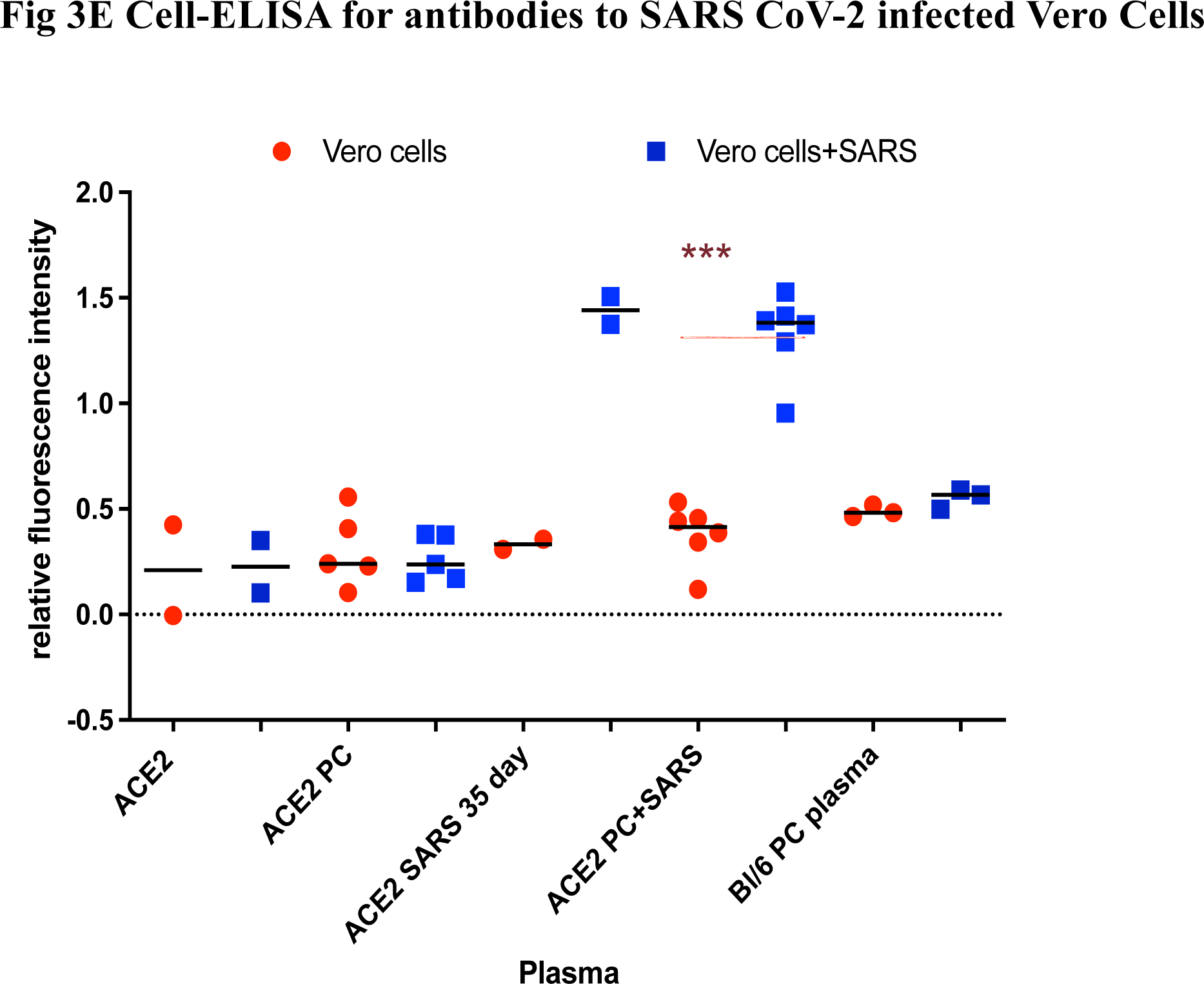
Survival, body temperature, SARS-CoV-2 viral load, cross reactivity of malarial antibodies with SARS- CoV-2 were monitored post challenge. Fig 3a) Kaplan–Meier analysis of percentage survival of control SARS-CoV-2 infected and K18 mice and animals prior recovered from malaria followed by challenge with SARS-CoV-2 are shown (*n* = 10 each). Fig 3b) Body temperature of infected mice of two groups was monitored daily for 20 days during the course of infection (*n* = 10 each). Fig 3c) Expression of nucleocapsid gene as by PCR in c-DNA of lungs of control SARS-CoV- 2 infected and recovered mice (N=2) and animals prior recovered from malaria followed by challenge with SARS-CoV-2 and recovered (N=6) are shown. Lungs of euthanized mice 35 days post challenge with SARS COV-2 were analysed by PCR Fig 3d) Demonstration of Plasmodia in circulation was determined by PCR with genus specific primers. DNA purified from blood were analysed on day 52 post viral challenge in mice recovered from malaria followed by challenge with SARS-CoV-2 are shown. Pre-SARS1 to Pre-SARS5 refers to blood collected from mice post recovery from malaria but before challenge with SARS CoV-2 PC+ SARS1-6 refers to PCR analysis of blood collected from animals recovered from SARS CoV-2 infection. Blood samples collected 35 days post challenge with are labelled as PC+SARS1-6. DNA purified from mice with acute Plasmodium chaubudi infection were taken as positive controls and are labelled as PC Positive 1 and 2. Fig 3e) Cell ELISA was performed to test serological cross-reactivity of malaria antibody with SARS-CoV-2 in plasma of different groups of mice. 96 well TC plates fixed with only Vero cells or Vero cells infected with SARS-CoV-2 were used for testing sera of mice. ACE2: Normal K18 ACE2 mice. ACE2 PC: K18 mice 30 days post Plasmodium chaubudi infection. ACE2 SARS 35 day: K18 mice recovered from SARS CoV-2 infection without prior exposure to malaria. ACE2 PC+SARS: K18 mice recovered from SARS CoV-2 infection with prior exposure to malaria (35 days). Bl/6 PC plasma: C57BL/6 mice 30 days post Plasmodium chaubudi infection.

### Effect of malaria exposure in chronic model of SARS-CoV-2 challenge infection

To address the issue of cross protection in terms of disease outcome to SARS-CoV-2 challenge infection by malaria, another cohort of mice were infected and followed up for mortality, changes in body temperature over a period of 35 days post challenge as described in the cartoon in Fig 1b. The logic was that this model would allow evaluation of *in vivo* cross-protection by malaria against SARS-CoV-2 in terms of severe pathology leading to mortality. The results shown in Fig 3a reveal statistically significant protection against mortality on challenge with SARS-CoV-2 in K18 mice that had experienced prior malaria infection – only 2 of the 9 controls survived while 6 of the 8 mice in the malaria pre-infected mice survived the challenge infection. Hypothermia, a characteristic feature of SARS- CoV-2 infection in K18 mice was observed in the control group on day 8 (Fig 3 b) in comparison to malaria recovered mice on challenge with SARS-CoV-2. Lung tissues of all the 8 recovered K18 mice were subjected to PCR and RT-PCR analysis 35 days post challenge. The results shown in Fig 3c and FigS3a indicate persistence of SARS-CoV-2 in both the control and malaria recovered mice. Analysis for persistence of *Plasmodium chabaudi* scored by PCR (16) in 6 of the Malaria – SARS-CoV- 2 recovered mice suggested that SARS-CoV-2 infection only marginally affects survival of malarial parasitemia in circulation (Fig 3d). These observations shown in Fig 3 c and <u>3d</u> suggest no significant adverse mutual impact on each other by SARS-CoV-2 and Plasmodium in co-infected animals at least during the period of the current study.

The possibility of antibodies generated by Plasmodial infections cross reacting with SARS-CoV-2 was tested by immunofluorescent cell based assay using Vero cells infected with the virus. Sera samples collected from mice post recovery from *Plasmodium chabaudi* infection (prior to challenge with SARS CoV-2) were tested. No significant cross-reactive antibodies could be detected. (Fig 3e). Sera collected from mice recovered from SARS-CoV-2 were used as positive controls which displayed high titers of antibodies to SARS-CoV-2 in the assay.

The current study provides definitive evidence to suggest that K18 hACE2 mice that had been exposed to malarial infections are protected against SARS-CoV-2 induced mortality although viral load in lungs were comparable between SARS-CoV-2 infected control mice in comparison to malaria co-infected animals when analyzed at one time point (6^th^ day) post challenge with SARS-CoV-2. In this context a recent report of Nippostrongyloides sp. infected K18 mice being resistant to subsequent challenge with virulent SARS-CoV-2 virus is highly significant - cross-protection by a nematode parasitic infection was demonstrable both in terms of decreased viral load in lungs as well as decreased mortality [21]. The study also provided evidence for the role of accumulation of CD8+ve cytotoxic T-cells generated by the developing worm stages in the lung contributing to decreased viral growth of SARS-CoV-2. This observation is more definitive of cross protection in mice that are co-infected with soil transmitted helminth infection than the current study in which no demonstrable anti-viral immunity (in terms of viral rolad in lungs) was observed in animals with past exposure to malaria. However, decreased mortality in malaria and SARS-CoV-2 co-infected mice is suggestive of development of functional resistance against mortality of SARS infected K18 mice. Inverse associations observed between malaria and COVID-19 in some of the epidemiological studies and attributing them to malaria (1 -6) would have to be interpreted with caution since several other tropical infections, particularly soil transmitted nematodes are highly prevalent in malaria endemic regions and may have contributed to lower morbidity and mortality due to Covid-19.

The conclusions drawn in the current study are not however free of caveats - K18 human ACE2 transgenic mice models are only surrogates that mimic COVID-19 in humans.

Although the model is very widely used for testing drugs and vaccine efficacy against challenge infections [21, 22] including co-infection studies, the model could be suboptimal to study physiological and immunological consequences of prior malaria exposure to understand mechanisms of cross protection against SARS CoV-2 infections. Human populations experience largely asymptomatic or mild low-grade self-limiting infections with SARS-CoV- 2 (17) but the murine K18 hACE2 transgenic mice model displays very high viral load and severe pathology when challenged with a single large bolus of virus leading to mortality of nearly 80-85% of mice. Similarly, self-limiting parasitaemia lasting about 15-20 days in murine malaria is different from malaria in human communities which are largely truncated by use of effective anti-malarial drugs. Regardless of these limitations, the current study can still be viewed as very significant since cross-protection to SARS-CoV-2 by malaria in terms of decreased mortality was demonstrable in the mouse model system of SARS-CoV-2 – a more severe infection model of disease in comparison to human COVID 19. The results of the current study do not offer any significant insights into precise mechanism of protection from mortality by prior exposure to malarial infections other than significant host responses in terms of difference in cytokine and chemokine induction in lungs and in circulation. However these differences cannot be causally attributed for the protection observed since it is not clear if these differential responses contributed significantly to survival of co-infected mice. We speculate that any or all of the following could have contributed for the observed protection against SARS-CoV-2 induced mortality: a) significant hematological disturbances caused by malarial infections in K18 hACE2 mice in terms of increased anemia, granulopoieses etc. <u>(Fig. S 4a-4f</u>) b) induction of trained immunity by long protracted malarial infections - Plasmodia are known to induce such immunity [23] or induction of tolerance that protects hosts against severe debilitating disease (18) c) significant changes in blood coagulation cascade that malaria are known to induce since adverse prognosis in COVID-19 is often attributed to blood coagulation in lungs [24] d) induction of auto-antibodies to carbohydrate moieties induced in malaria [20, 25] that have been proposed to play a role in COVID-19. The study provides unambiguous evidence for absence of anti-viral antibody mediated cross protection in terms of viral multiplication in lungs as a consequence of co-infection with malaria. Epidemiological investigations on co-infections with two or more pathogens in human and animal communities (including influence of Malaria and SARS-CoV-2) are limited by known and several unknown confounding factors and experimental studies in animal models with all inherent limitations are the only available recourse for validation of such associations.

## Supporting information

Fif Sup

## Acknowledgements

The Institute of Life Sciences, Bhubaneswar is a constituent Institute fully funded by the Department of Biotechnology, Ministry of Science and Technology, Government of India. The core facility of ABSL3 laboratory was funded by BIRAC operating under DBT and by a grant number: BT/PR44372/MED/29/1580/2021 from Department of Biotechnology, Government of India.

## References

1. Chen, C., et al., Global prevalence of post-coronavirus disease 2019 (COVID-19) condition or long COVID: a meta-analysis and systematic review. The Journal of infectious diseases, 2022. 226(9): p. 1593–1607.

2. Anyanwu, M.U., The association between malaria prevalence and COVID-19 mortality. BMC Infectious Diseases, 2021. 21(1): p. 1–6.

3. Hussein, M.I.H., et al., Malaria and COVID-19: unmasking their ties. Malaria journal, 2020. 19(1): p. 1–10.

4. Orish, V.N., et al., Is malaria immunity a possible protection against severe symptoms and outcomes of COVID-19? Ghana medical journal, 2021. 55(2): p. 56–63.

5. Rusmini, M., et al., How genetics might explain the unusual link between malaria and COVID-19. Frontiers in Medicine, 2021. 8: p. 650231.

6. Mahajan, N.N., et al., Co-infection of malaria and early clearance of SARS-CoV-2 in healthcare workers. Journal of medical virology, 2021. 93(4): p. 2431–2438.

7. Wildner, N.H., et al., B cell analysis in SARS-CoV-2 versus malaria: increased frequencies of plasmablasts and atypical memory B cells in COVID-19. Journal of Leucocyte Biology, 2021. 109(1): p. 77–90.

8. Iesa, M., et al., SARS-CoV-2 and Plasmodium falciparum common immunodominant regions may explain low COVID-19 incidence in the malaria-endemic belt. New microbes and new infections, 2020. 38: p. 100817.

9. Manning, J., et al., SARS-CoV-2 cross-reactivity in prepandemic serum from rural malaria-infected persons, Cambodia. Emerging infectious diseases, 2022. 28(2): p. 440.

10. Suresh, V., et al., Quantitative proteomics of hamster lung tissues infected with SARS-CoV-2 reveal host factors having implication in the disease pathogenesis and severity. The FASEB Journal, 2021. 35(7).

11. Arce, V.M. and J.A. Costoya, SARS-CoV-2 infection in K18-ACE2 transgenic mice replicates human pulmonary disease in COVID-19. Cellular & Molecular Immunology, 2021. 18(3): p. 513–514.

12. Winkler, E.S., et al., SARS-CoV-2 infection of human ACE2-transgenic mice causes severe lung inflammation and impaired function. Nature immunology, 2020. 21(11): p. 1327–1335.

13. Guha, S.K., et al., Single episode of mild murine malaria induces neuroinflammation, alters microglial profile, impairs adult neurogenesis, and causes deficits in social and anxiety-like behavior. Brain, behavior, and immunity, 2014. 42: p. 123–137.

14. Singh, B., et al., Isolation and Characterization of Five Severe Acute Respiratory Syndrome Coronavirus 2 Strains of Different Clades and Lineages Circulating in Eastern India. Frontiers in microbiology, 2022. 13: p. 856913.

15. Mendoza, E.J., et al., Two Detailed Plaque Assay Protocols for the Quantification of Infectious SARS-CoV-2. Curr Protoc Microbiol, 2020. 57(1): p. ecpmc105.

16. Perkins, et al., Use of PCR for detection of subpatent infections of lizard malaria: implications for epizootiology. Molecular Ecology, 1998. 7(11): p. 1587–1590.

17. Chen I et al (2016) “Asymptomatic” Malaria: A chronic and debilitating Infection that should be treated. PLoS Med 13(1): e1001942

18. Nahrendorf., et al,.Inducible mechanisms of disease tolerance provide an alternative strategy of acquired immunity to malaria eLife 2021;10:e63838. DOI: 10.7554/eLife.63838

19. Zhong, J., et al., Robust hepatitis C virus infection in vitro. Proceedings of the National Academy of Sciences, 2005. 102(26): p. 9294–9299.

20. Lapidus, S., et al., Plasmodium infection induces cross-reactive antibodies to carbohydrate epitopes on the SARS-CoV-2 Spike protein. MedRxiv, 2021.

21. Oyesola, O.O., et al., Exposure to lung-migrating helminth protects against murine SARS-CoV-2 infection through macrophage-dependent T cell activation. Science Immunology, 2023. 8(86): p. eadf8161.

22. Dong, W., et al., The K18-human ACE2 transgenic mouse model recapitulates non-severe and severe COVID-19 in response to an infectious dose of the SARS-CoV-2 virus. Journal of virology, 2022. 96(1): p. e00964–21.

23. Netea, M.G., et al., The role of trained immunity in COVID-19: Lessons for the next pandemic. Cell Host & Microbe, 2023. 31(6): p. 890–901.

24. Mackman, N., et al., Coagulation abnormalities and thrombosis in patients infected with SARS-CoV-2 and other pandemic viruses. Arteriosclerosis, thrombosis, and vascular biology, 2020. 40(9): p. 2033–2044.

25. Breiman, A., et al., Low levels of natural anti-α-N-acetylgalactosamine (Tn) antibodies are associated with COVID-19. Frontiers in Microbiology, 2021. 12: p. 141.

