## Supplementary material for "Prior exposure to malaria decreases SARS-CoV-2 mediated mortality in K18-hACE2 mice without influencing viral load in lungs": Fif Sup

**Sup File Mawatwal et al**

**Supplementary Figures and Legends:-**

**Fig S1.**

**Suplementary Fig S1**

**Fig. S1.** **Expression of cytokines, chemokines and IFN-β in lung at 6 d.p.i**

**a.**

**b.**

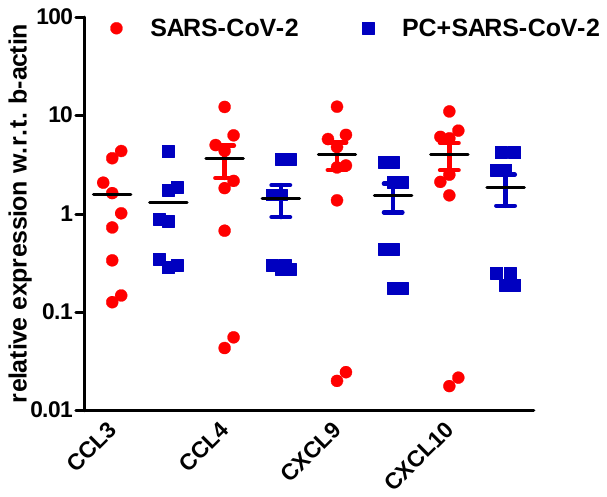

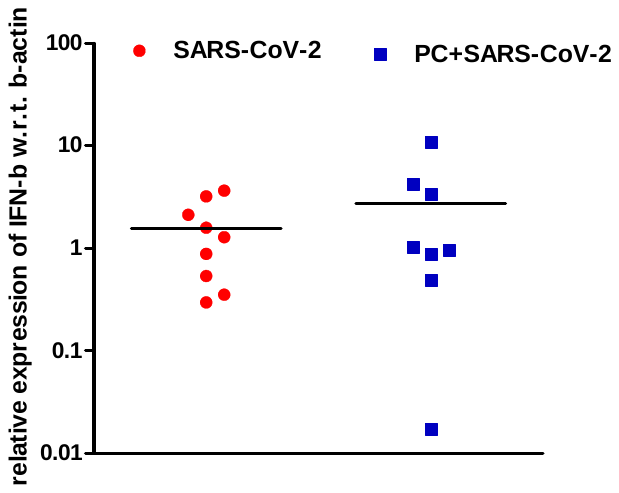

**d**.

**c.**

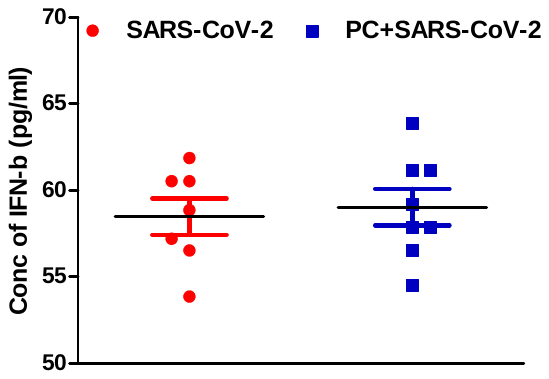

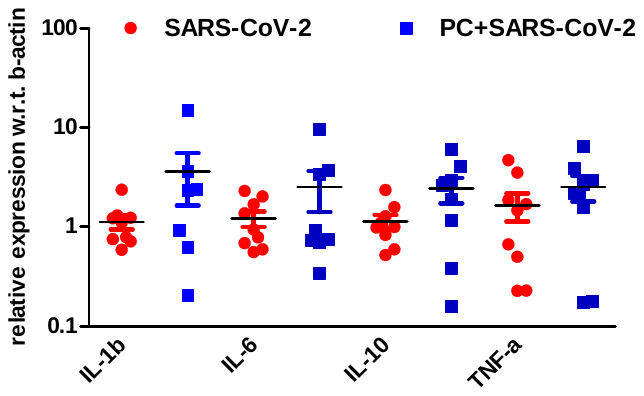

**Legends for Supplementary FigS1**

**a)** qRT-PCR analysis for indicated chemokines with total RNA isolated from lungs of only SARS-CoV-2 infected (n = 9) and Malaria + SARS-CoV-2 infected (n = 8) ACE2 mice at 6 d.p.i. Data are presented as mean ± SEM. Statistical analyses were performed by Student’s unpaired t-test.

**b)** qRT-PCR analysis for Interferon-b using mRNA isolated from lungs of only SARS CoV2 infected (n = 9) and PC + SARS CoV2 infected (n = 8) ACE2 mice at 6 d.p.i. Data are presented as mean ± SEM. Statistical analyses were performed by Student’s unpaired t-test.

**c)** Plasma levels of IFN- β measured by ELISA only SARS-CoV-2 infected (n = 9) and Malaria + SARS-CoV-2 infected (n = 8) ACE2 mice at 6 d.p.i. Data are presented as mean ± SEM in pg/ml of plasma.

**d)** Cytokine expression in in the lungs of only SARS-CoV-2 infected (n = 9) and Malaria + SARS-CoV-2 infected (n = 8) ACE2 mice at 6 d.p.i. was determined by qRT-PCR. Data are presented as mean ± SEM. Statistical analyses were performed by Student’s unpaired t-test.

**Fig S2. Expression of SARS-CoV-2 nucleocapsid in lungs at 35 and 6 d.p.i :**

b.

a.

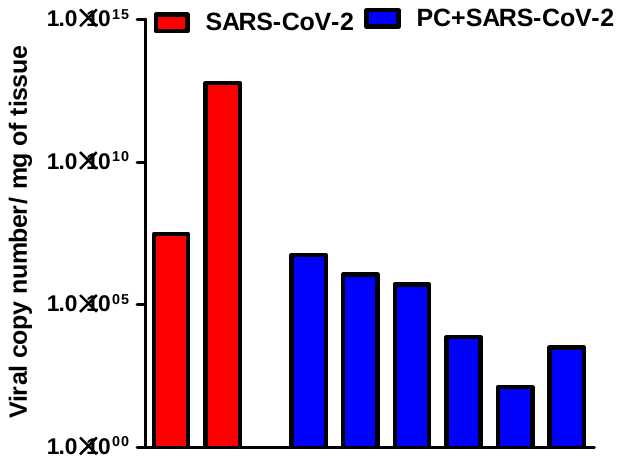

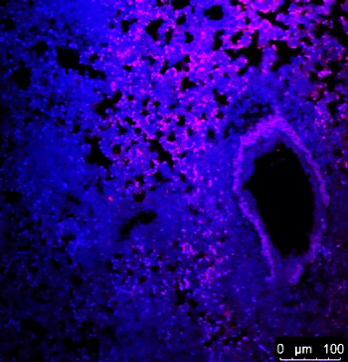

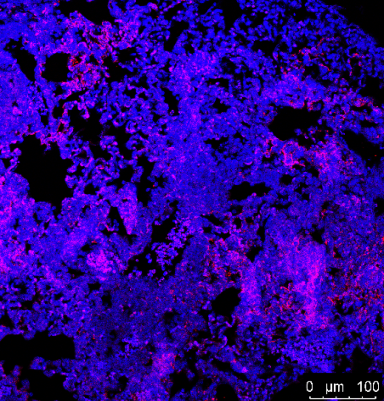

SARS-CoV-2	PC+SARS-CoV-2

c.

SARS-CoV-2	PC+SARS-CoV-2

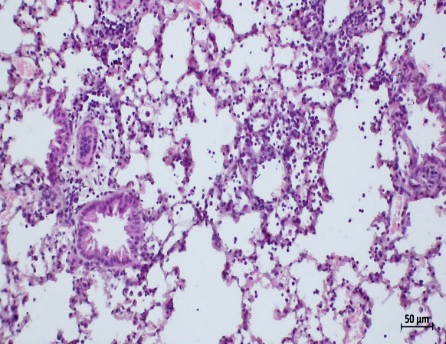

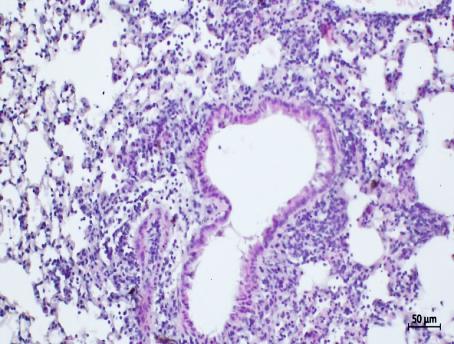

d.

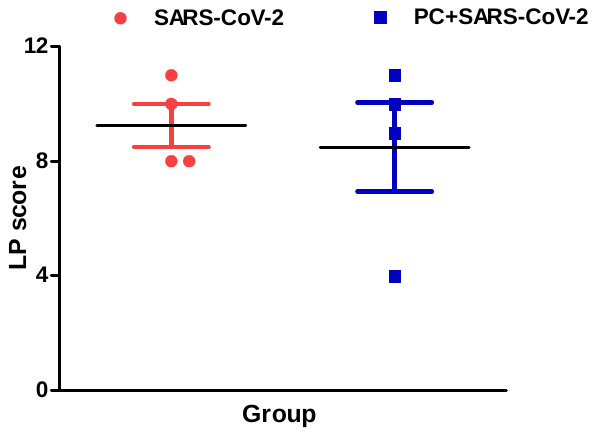

**Legends Fig S2:**

**a)** qRT-PCR SARS-CoV-2 Viral copy number analysis in lungs of K18 mice infected only with SARS CoV2 (red bars) or Malaria + SARS-CoV-2 group in ACE2 mice 35 days post SARS infection. Data shown is only for each of the surviving mice in both the groups

**b)** Representative immunofluorescent microscopy images of lung sections probed with antibodies to SARS-CoV-2 nucleocapsid (NP). Left: lungs of a K18 mouse infected only with SARS-CoV-2. Right: lungs of a K18 mouse infected with SARS CoV-2 with prior exposure to Plasmodium infectin. Lungs collected 6 days post challenge with SARS COV-2 were sectioned for analysis.

**c)** Hematoxylin and eosin-stained lung sections of same animals shown above in FigS3 b above.

**d)** Lung pathology score in lung lysates of only SARS-CoV-2 infected and PC + SARS CoV2 infected ACE2 mice at 6 d.p.i. Data are presented as mean ± SEM. Statistical analyses were performed by Student’s unpaired t-test.

**Fig S3 Parasitaemia and heamatological changes in C57BL/6 mice and K18 hACE2 Tg mice**

b.

a.

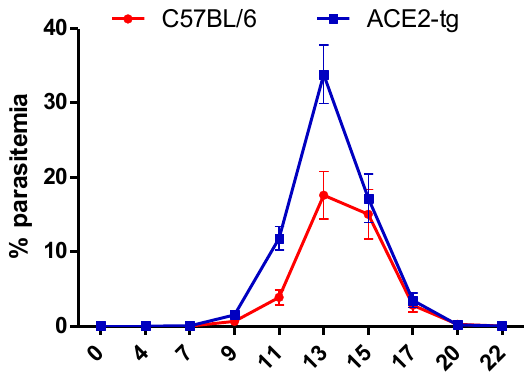

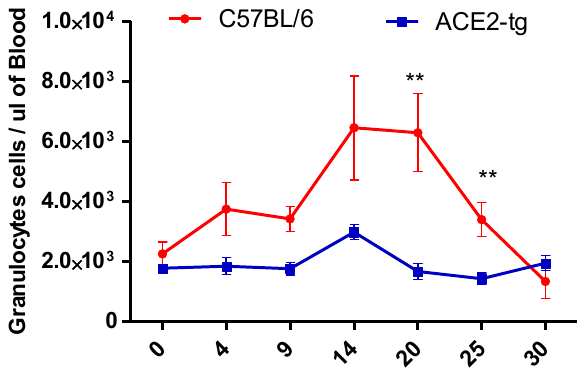

d.

c.

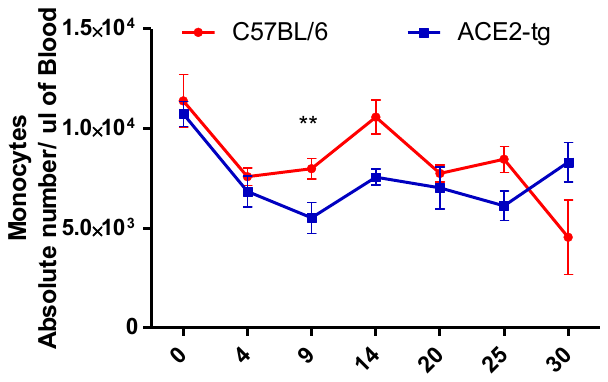

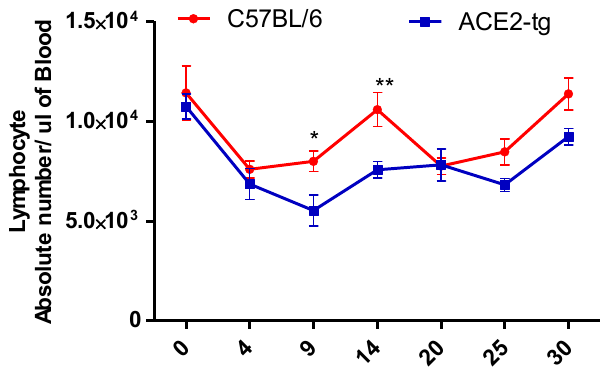

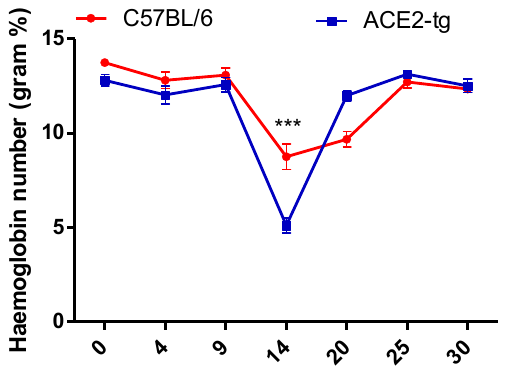

e.

**Legends for Fig S3: Parasitaemia and heamatological changes in C57BL/6 mice and K18 hACE2 Tg mice infected with Plasmodium chaubudi :**

a) Parasiatemia in C57BL/6 and ACE2- transgenic mice blood b) absolute granulocyte count c) absolute monocytes d) absolute lymphocytes and e) haemoglobin (Mean SEM of n=10). Statistical significance is marked in Asterix.
